## Appendix for "CellPLM: Pre-training of Cell Language Model Beyond Single Cells"

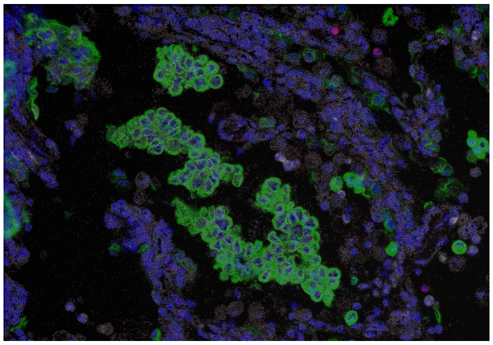

(a) Visualization of molecular image.

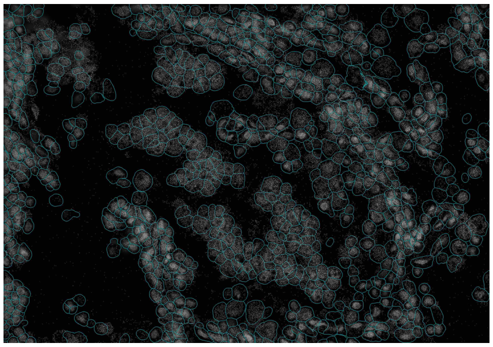

(b) Visualization of cell segmentation.

Figure 4: (a) A sample image of protein and RNA molecules. (b) A sample image of segmented cells.

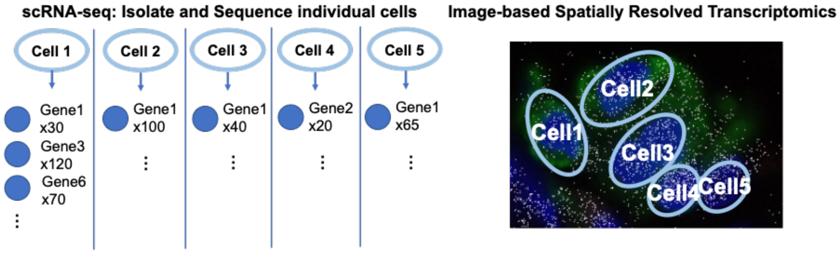


Figure 5: An illustration of the difference between scRNA-seq and SRT data.

#### B 2D Sinusoid Positional Encodings

Since 2D sinusoidal PE achieves a competitive performance and has a lower complexity on SRT data Wen et al. (2023), in our transformer encoer, we generate a sinusoidal PE for cells in SRT data, formulated as:

$$\begin{aligned}
&\text{PE}_{(x,y,2i)} = \sin\left(x/10000^{4i/d}\right), \text{PE}_{(x,y,2i+1)} = \cos\left(x/10000^{4i/d}\right), \\
&\text{PE}_{(x,y,2j+d/2)} = \sin\left(y/10000^{4j/d}\right), \text{PE}_{(x,y,2j+1+d/2)} = \cos\left(y/10000^{4j/d}\right),
\end{aligned} \tag{8}$$

where $d$ is the total dimension of positional encoding, $i, j \in [0, d/4)$ specify a specific feature dimension. Let $\tilde{\mathbf{C}} \in \mathcal{R}^{N\times 2}$ be a normalized coordinate matrix, where we normalize and truncate coordinates in $\mathbf{C}$ to integers ranging in $[0, 100)$. $x, y$ then refer to the spatial coordinates from $\tilde{\mathbf{C}}$, e.g., $x = \tilde{\mathbf{C}}_{t,0}$ and $y = \tilde{\mathbf{C}}_{t,1}$ for cell $t$. In this way, we generate a PE matrix $\mathbf{P} \in \mathcal{R}^{N\times d}$ for every cell in SRT data, where $\mathbf{P}_i$ is the PE vector for cell $i$. Meanwhile, for scRNA-seq data, a randomly initialized $d$-dimensional vector $p'$ is shared among all cells, which also results in a placeholder PE matrix $\mathbf{P}$.

$$\begin{aligned} p(\mathbf{y}_i; \boldsymbol{\pi}) &= \text{Multinomial}(\boldsymbol{\pi}), \\ p(\mathbf{z}_i \mid \mathbf{y}_i) &= \prod_{i=1}^{L} \mathcal{N}\left(\boldsymbol{\mu}_{y_{i,l}}, \text{diag}\left(\boldsymbol{\sigma}^2_{y_{i,l}}\right)\right), \\ p_{\theta_{dec}}(\mathbf{x}_i \mid \mathbf{z}_i) &= \mathcal{N}\left(\boldsymbol{\mu}_{\mathbf{z}_i}, \sigma^2 \mathbf{I}\right), \end{aligned} \tag{9}$$

where $\mathbf{y}_i \in \mathcal{R}^L$ represents the one-hot latent cluster variable and $\boldsymbol{\pi}$ is its prior; $y_{i,l}$ denotes the $l$-th entry of $\mathbf{y}_i$; $\boldsymbol{\mu}_{y_l} \in \mathcal{R}^{d_z}$ and $\boldsymbol{\sigma}^2_{y_l} \in \mathcal{R}^{d_z \times d_z}$ denote the mean and variance of the $l$-th Gaussian component, respectively; and $\boldsymbol{\mu}_{z_i} \in \mathcal{R}^k$ and $\sigma^2 \mathbf{I} \in \mathcal{R}^{k \times k}$ denote the posterior mean and variance of expression $\mathbf{x}_i$, respectively. In this work, we assume that $\sigma^2$ is a constant and the posterior mean is parameterized by $\boldsymbol{\mu}_{z_i} = f_{dec}(\mathbf{z}_i; \theta_{dec})$.

To estimate the posterior of $\mathbf{z}_i$ and $\mathbf{y}_i$, we parameterize the inference process with neural networks. Specifically, we assume that the cluster variables $\mathbf{y}$ are independent of the expression $\mathbf{x}_i$ condition on latent variables $\mathbf{z}_i$. The inference model can be formulated as:

$$\begin{aligned} q_{\eta_\mu, \eta_\sigma}(\mathbf{z}_i \mid \mathbf{x}_i) &= \mathcal{N}\left(\hat{\boldsymbol{\mu}}_i, \text{diag}\left(\hat{\boldsymbol{\sigma}}^2_i\right)\right), \\ q_{\eta_\pi}(\mathbf{y}_i \mid \mathbf{z}_i) &= \text{Multinomial}(\hat{\boldsymbol{\pi}}_i), \end{aligned} \tag{10}$$

where the estimations are given by

$$\begin{aligned} \mathbf{h}_i &= f_{enc}(\mathbf{x}_i; \eta_{enc}), \\ \hat{\boldsymbol{\mu}}_i &= f_\mu(\mathbf{h}_i; \eta_\mu), \\ \log\left(\hat{\boldsymbol{\sigma}}^2_i\right) &= f_\sigma(\mathbf{h}_i; \eta_\sigma), \\ \hat{\boldsymbol{\pi}}_i &= f_\pi(\mathbf{z}_i; \eta_\pi). \end{aligned} \tag{11}$$

Here $f_{enc}(\cdot; \eta_{enc})$ represents the transformer encoder, $f_\mu(\cdot; \eta_\mu)$, $f_\sigma(\cdot; \eta_\sigma)$ and $f_\pi(\cdot; \eta_\pi)$ are neural networks. A log-evidence lower bound (ELBO) can be derived from this generative model for the optimization purpose (Dilokthanakul et al., 2016). However, as mentioned in Section 3.1, our pre-training framework incorporates a cell language model, where parts of the input gene expression matrix $\mathbf{X}$ are masked. This will result in a modified objective. To formalize the problem, recall that previously we defined the masked set as $\mathcal{M}$. On top of that, we denote $\mathbf{M} \in \mathcal{R}^{N \times k}$ as a mask indicator matrix such that

$$\mathbf{M}_{i,j} = \begin{cases} 1 & \text{if } (i,j) \notin \mathcal{M}, \\ 0 & \text{if } (i,j) \in \mathcal{M}. \end{cases}$$

Let $\tilde{\mathbf{X}} \in \mathcal{R}^{N \times k}$ be the masked gene expression matrix given by the element-wise multiplication $\tilde{\mathbf{X}} = \mathbf{M} \odot \mathbf{X}$. The objective of cell language model with Gaussian mixture prior, i.e., a denoising variational lower bound (Im Im et al., 2017), can be formulated as:

$$\begin{aligned} \mathcal{L}_{\text{CellLM}} =& \mathbb{E}_{q(\mathbf{Z},\mathbf{Y}|\tilde{\mathbf{X}})} \mathbb{E}_{p(\tilde{\mathbf{X}}|\mathbf{X})} \left[ \ln \frac{p_\theta(\mathbf{X}, \mathbf{Z}, \mathbf{Y})}{q_\eta(\mathbf{Z}, \mathbf{Y} \mid \tilde{\mathbf{X}})} \right] \\ =& \underbrace{\mathbb{E}_{q_{\eta_{enc}}(\mathbf{Z}|\tilde{\mathbf{X}})} \mathbb{E}_{p(\tilde{\mathbf{X}}|\mathbf{X})} \left[ \log p_{\theta_{dec}}(\mathbf{X} \mid \mathbf{Z}) \right]}_{\mathcal{L}_{\text{recon}}} - \underbrace{\mathbb{E}_{q_{\eta_\pi}(\mathbf{Y}|\mathbf{Z})} \left[ \text{KL}\left( q_{\eta_{enc}}(\mathbf{Z} \mid \tilde{\mathbf{X}}) \| p(\mathbf{Z} \mid \mathbf{Y}) \right) \right]}_{\mathcal{L}_{\text{cond}}} \\ & - \underbrace{\mathbb{E}_{q_{\eta_{enc}}(\mathbf{Z}|\tilde{\mathbf{X}})} \left[ \text{KL}\left( q_{\eta_\pi}(\mathbf{Y} \mid \mathbf{Z}) \| p(\mathbf{Y}) \right) \right]}_{\mathcal{L}_{\text{Y}}}. \end{aligned} \tag{12}$$

### E Pre-training Settings

#### E.1 Hyperparameter Settings

We pre-trained *CellPLM* model with the hyperparameters specified in Table 5.

| | *CellPLM* |
|---|---|
| encoder hidden dim | 1024 |
| encoder layers | 4 |
| latent dimension | 512 |
| decoder hidden dim | 1024 |
| decoder layers | 2 |
| model dropout | 0.2 |
| cell mask rate | 0.75 |
| gene mask rate | 0.25 |
| learning rate | 2e-4 |
| weight decay | 1e-8 |
| num of cluster (for GMM) | 16 |
| total parameter | 82,402,543 |

Table 5: Hyperparameters for pretraining *CellPLM* model.

##### E.2 Datasets for Pre-training

The dataset for pre-training contains $11.4$ million cells from scRNA-seq and SRT data. scRNA-seq data consist of $4.7$ million cells from human tumor cell atlas (HTCA, `https://humantumoratlas.org/`), $1.4$ million cells from human cell atlas (HCA, `https://www.humancellatlas.org/`), and $2.6$ million cells from Gene Expression Omnibus (GEO, `https://www.ncbi.nlm.nih.gov/geo/`). All of them are public available data. A more detailed list will be disclosed in our GitHub repository. Note that although our *CellPLM* is capable to handle various input feature sets, when we concatenated these scRNA-seq datasets, we used inner join by default of Anndata package. As a result, all scRNA-seq datasets only contain a $13,500$ common gene set. We will address this issue and increase the size of the gene set in future versions of *CellPLM*.

The SRT datasets we used are publicly available on Nanostring official website: `https://nanostring.com/products/cosmx-spatial-molecular-imager/nsclc-ffpe-dataset/`, where $2.7$ million cells and $1,000$ genes are measured.

#### F Additional Experimental Details

In this section, we provide more experimental details about fine-tuning, baselines, and evaluation metrics under each downstream task.

##### F.1 scRNA-seq Denoising

**Downstream Task Datasets.** In scRNA-seq denoising task, we evaluate *CellPLM* on two datasets, i.e., PBMC 5K and Jurkat from 10x Genomics lin (a). It is worth noting that during the prepossessing stage, we performed sub-setting on both datasets to ensure that all the genes were included in the gene set of pre-training data. Additionally, any genes with zero counts were removed from the analysis. We list the statistics of them in Table 6.

Table 6: scRNA-seq denoising datasets

| | 5K PBMC | Jurkat |
|---|---|---|
| Number of genes | 33,538 | 32,738 |
| Number of cells | 5,247 | 3,258 |
| Num genes picked | 7,197 | 7,618 |

**Evaluation Metrics.** Following the setting of scGNN Wang et al. (2021), scGNN2.0 Gu et al. (2022) and DeepImpute Arisdakessian et al. (2019), we performed synthetic dropout simulation with missing at random (MAR) setting. While scGNN only considered a simple scenario, i.e., randomly flipped $10\%$ of the non-zero entries to zeros, DeepImpute applied cell-wise mask with masking probability given by a multinomial distribution. Specifically, we adapted the setting from DeepImpute with exponential kernel. For cell $i$ that contains at least $5$ expressed genes, the probability that one non-zero count $x_{i,j}$ is masked during the training process is given by $\text{Exp}(0, 20)$:

$$p_{i,j} = \frac{1}{20} e^{-\frac{x}{20}},$$
$$q_{i,j} = \frac{p_{i,j}}{\sum_{j=0}^{J_i} p_{i,j}},$$

where $J_i$ is the number of non-zero counts within cell $i$. We masked $10\%$ of the non-zero counts according to $\{q_{i,j}\}_{j=0}^{J_i}$ and evaluate model performance on the masked entries. We calculate the root mean squared error (RMSE) and mean absolute error (MAE) between the predicted values and ground truth.

Table 7: Spatial tanscriptomic imputation datasets.

| | Lung2 | Liver2 | GSE131907 | GSE151530 |
|---|---|---|---|---|
| Number of genes | 500 | 500 | 29,634 | 18,667 |
| Number of cells | 836,739 | 598,141 | 208,506 | 56,721 |
| Num genes picked | 462 | 446 | All | ALL |
| Num cells picked | 40,114 | 20,629 | All | All |

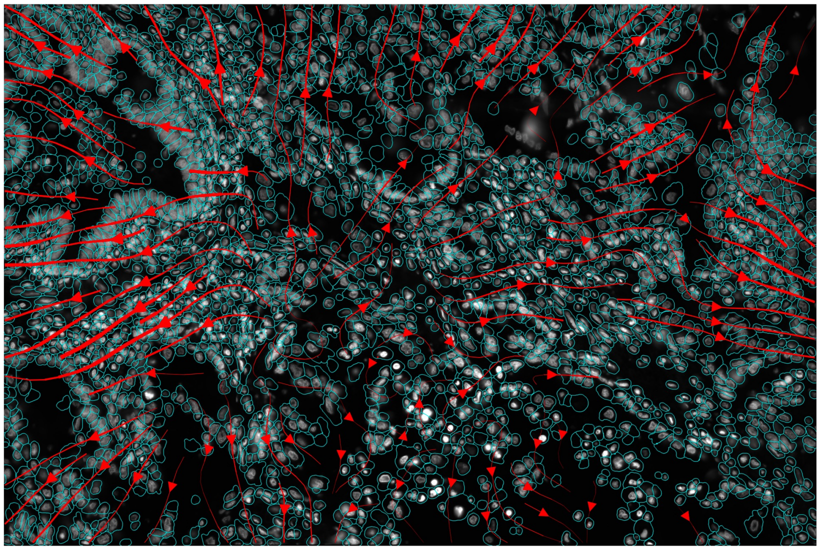

Figure 6: Visualization of attention matrix demonstrate cell-cell communication.

**Visualization of attention.** One essential multi-cell task is cell-cell communication (CCC) inference, where CCC mainly represents biochemical signaling through ligand-receptor binding across cells (Cang et al., 2023). Our *CellPLM* applies self-attention mechanism on cell level, from which we can study the interaction strength given by cell attention matrix. As a preliminary study, we extract the attention matrix between cells from a random chosen field of view (FOV) in Cosmx Liver dataset. The attention matrix is treated as CCC scores, and we visualize the results following the stream plot setting in Cang et al. (2023). As shown in the Figure 6 in our supplementary PDF, there are some strong trends on the left side and right side of the FOV, suggesting further exploration of specific signaling pathways for the included cells. This case study showcase the potential of our *CellPLM* model in cell-cell communication research. We hope our model can facilitate more insightful biological research in the future.

##### F.3 Perturbation Prediction

The perturb-seq technology has been established to examine the gene expression response at the single-cell level when subjected to pooled perturbations (Dixit et al., 2016). By comparing the gene expression before and after perturbation, downstream analysis of differential expression (DE) enables the identification of genes that play a crucial role in disease progression. To assess the potential benefits of *CellPLM* in gene-level tasks, we conduct experiments to predict the expression value of genes after perturbation. Following the setting of GEARS (Roohani et al., 2022), we partition the perturbations into training, validation, and test sets, ensuring that none of the test perturbations are encountered during the optimization process.

Two perturbation datasets are employed for evaluation: (1) the Adamson Perturb-Seq dataset (Adamson et al., 2016), consisting of 87 one-gene perturbations; and (2) the Norman Perturb-Seq dataset (Norman et al., 2019), containing 131 two-gene perturbations and 105 one-gene perturbations. To evaluate the performance of perturbation prediction, we employ Root Mean Square Error (RMSE) to measure the degree of similarity between the non-zero ground-truth expression values and corresponding predicted gene expressions. In addition, following previous settings in GEARS (Roohani et al., 2022), we also present the RMSE calculated on the top 20 deferentially-expressed genes.

We compare the performance between *CellPLM* and two baselines, i.e., a recent preprint GEARS method (Roohani et al., 2022), and scGen (Lotfollahi et al., 2019). The results in Figure 7 imply that *CellPLM* achieves the lowest RMSE values across all settings.

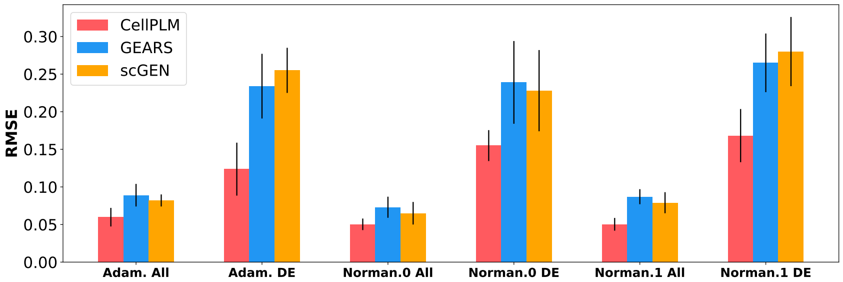


Figure 7: (*Task 3*) The RMSE performance (↓) on Adamson Perturb-Seq and the Norman Perturb-Seq datasets. The Norman Perturb-seq dataset consists of two settings: one-gene perturbations and two-gene perturbations, denoted as Norm.0 and Norm.1, respectively.

**Downstream Task Datasets.** We included the Adamson Perturb-Seq dataset Adamson et al. (2016) for one-gene perturbations and the Norman Perturb-Seq dataset Norman et al. (2019) for two-gene perturbations. We followed the preprocess pipeline of GEARS Roohani et al. (2022) and both datasets were then gene-wise subsetted to fit in the gene set of pre-training data. The statistics are summaried in Table 8.

Table 8: Perturbation prediction datasets.

| | Adamson | Norman |
|---|---|---|
| Number of genes | 5,060 | 5,045 |
| Number of cells | 68,603 | 91,205 |
| Num genes picked | 3,246 | 2,353 |
| Num one-gene pert. | 87 | 105 |
| Num two-gene pert. | – | 131 |

**Evaluation Metrics.** Following the setting of GEARS Roohani et al. (2022), we applied data split such that the testing perturbation are unseen during the training process. Specifically, For Adamson dataset, we randomly hold out 25% of the perturbations for testing and 10% of the perturbations within the training set for validation. For Norman dataset, two settings for two-gene perturbations are implemented for evalutation purpose: $1/2$ unseen and $2/2$ unseen. We excluded all two-gene combinations in which at least one of the individual genes involved in the combination belonged to the unseen set. Finally, we evaluate the performance by calculating the root mean squared error (RMSE) between the predictions and the true values within the testing set.
